# Mass spectrometry-based proteomics delivers in-depth proteome profiling of FFPE lung cancer biopsies from single glass slides

**DOI:** 10.64898/2026.01.20.700495

**Authors:** Olena Berkovska, Igor Schliemann, Georgios Mermelekas, Nazlı Ezgi Özkan, Mahnaz Nikpour, Vilde Drageset Haakensen, Åslaug Helland, Janne Lehtiö, Lukas M. Orre

## Abstract

Clinical proteomics has the potential to add a valuable data layer to genomic and histopathological analyses in precision oncology, but its application to limited clinical material remains challenging. Here, we demonstrate that state-of-the-art mass spectrometry-based proteomics enables in-depth proteomic profiling of formalin-fixed paraffin-embedded (FFPE), including small biopsies and single tissue sections mounted on glass slides. Despite minimal input material, single-slide analyses of clinical lung tumor specimens yielded biologically and clinically informative data, supporting detection of actionable proteins, immune-related signatures, and multivariate biomarkers. These results establish the feasibility of proteomics for retrospective FFPE studies and routine clinical practice, expanding opportunities for biomarker discovery and precision medicine from scarce tissue material.

## Main text

Proteomics has emerged as a promising complement to immunohistochemistry (IHC) and genomic profiling in cancer research and diagnostics, offering deeper insight into tumor biology and therapeutic response^1-6^. To date, most proteomics studies have relied on fresh-frozen tissue; however, formalin-fixed paraffin-embedded (FFPE) samples are the most widely used material in clinical practice, with millions of specimens stored in biobanks. FFPE material therefore holds great potential for the large-scale studies needed to decipher cancer complexity and advance precision medicine. Substantial progress has been made in mass spectrometry (MS)-based proteomics of FFPE samples, driven by advances in instrument sensitivity and improved workflows^7,8^. Nonetheless, performance in clinical settings with limited material, such as biopsies and/or tissue sections mounted on glass slides, remains unclear.

In this study, we analyzed a cohort of FFPE tumor biopsy specimens from unresectable lung cancer using a single 4-μm section on a glass slide per patient (“single-slide cohort”, n = 68; **Figure 1A**; **Supplementary Table 1**). An additional, smaller cohort comprising multiple FFPE scrolls from biopsies or surgically resected tumor specimens served as a reference (“multi-section cohort”, n = 15). The samples were analyzed using label-free data-independent acquisition (DIA) proteomics on two state-of-the-art MS instruments (Orbitrap Astral and timsTOF HT). In addition, multi-section cohort samples were analyzed on an older-generation MS instrument (Orbitrap Exploris).

**Figure 1.**
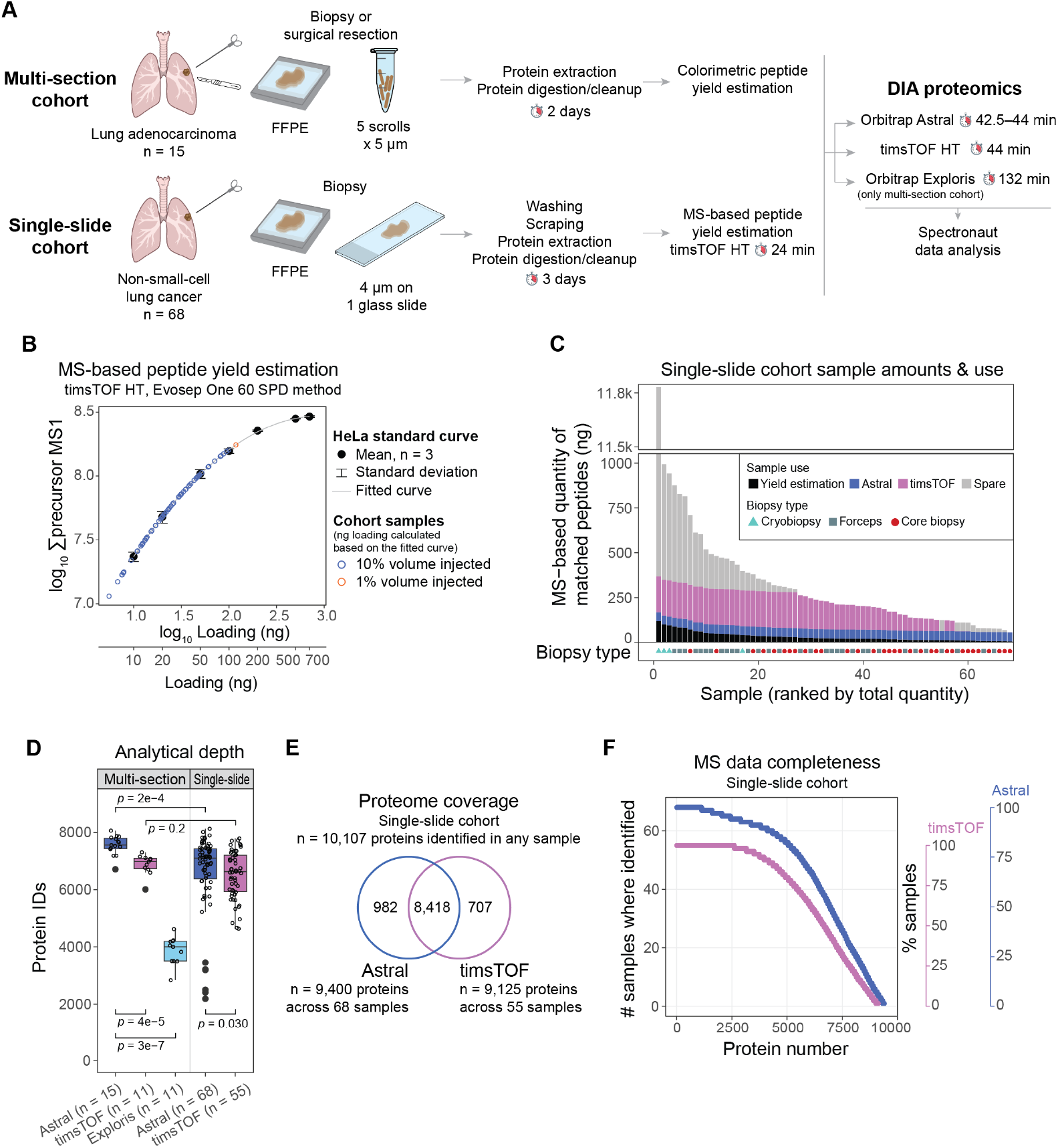
MS-based analysis of formalin-fixed, paraffin embedded (FFPE) clinical tumor samples delivers in-depth, proteome-wide profiling. **A)** Schematic overview of the study design. **B)** HeLa-based standard curve relating injected peptide amount to total MS1 signal, with single-slide cohort samples (n = 68) projected onto the curve to estimate injected peptide amounts. The curve was modeled using a quadratic polynomial regression relating log_10_-transformed values. **C)** Estimated peptide yields extracted from glass slides in the single-slide cohort (calculated as shown in panel B) and the proportion of each sample used for the different analyses. **D)** Number of proteins (gene-centric) identified per sample in MS-based analyses of the multi-section and single-slide cohorts. *P* values were calculated using two-sided Wilcoxon rank-sum test. The number of samples in each dataset is indicated in the figure. **E)** Total number of proteins (gene-centric) identified in the single-slide cohort analyses and the overlap between the Astral and timsTOF datasets. **F)** Number of samples in which each protein was identified in the single-slide cohort analyses.

Sample preparation for both cohorts followed the same general workflow, with two key differences: inclusion of a wash step for glass slides and the use of an MS-based peptide amount estimation instead of a colorimetric assay (**Figure 1A**). In a small pilot analysis (n = 6) of the single-slide cohort, we observed substantially higher MS signal from skin keratins compared with the multi-section cohort (**Figure S1A,B**). High-abundance proteins present a challenge in MS proteomics due to ion suppression of lower-abundance peptide signals. We hypothesized that the elevated skin keratin signal resulted from contamination accumulated during handling and long-term storage of glass slides. Accordingly, we introduced a glass-washing step using a mild non-ionic detergent for preparation of the full cohort. Although this step modestly reduced skin keratin levels, the difference was not significant, and the proportion of MS signal originating from a small number of high-abundance proteins remained significantly higher than in the multi-section cohort (**Figure S1B**). Further protocol optimization may improve these results but was beyond the scope of this study due to limited material availability. Based on our observations, we recommend handling slides with gloves and using freshly sectioned samples or storing slides in sealed containers.

The second protocol adaptation for the single-slide cohort involved peptide yield estimation. Colorimetric assays are commonly used in bulk proteomics to quantify total peptide yield and normalize injection amounts; however, they require relatively large amounts of material (several hundred nanograms), and their accuracy can be compromised by interference from non-peptide components in FFPE samples. To address these limitations, we employed an MS-based estimation strategy similar to that described by Tüshaus *et al*.^7^ Briefly, single-slide cohort samples were analyzed using a fixed injection volume alongside HeLa standards (10–700 ng injections). A HeLa standard curve was generated using summed MS1 precursor signal, and peptide amounts in clinical samples were calculated from this curve (**Figure 1B**). We refer to these values as “MS-based quantity of matched peptides”, as this estimate does not account for fragmented, cross-linked or otherwise non-canonical peptides.

MS-based matched peptide yields in the single-slide cohort ranged from 55 ng to 1 μg per sample, except for one large cryobiopsy yielding more than 10 μg (**Figure 1C**). These values were substantially lower than those obtained from the multi-section cohort, in which 5–250 μg of colorimetrically quantified peptides were extracted (**Figure S1C**). Notably, biopsy size showed a moderate-to-strong correlation with the estimated peptide yield (**Figure S1D**). Although biopsy length cannot be used to reliably normalize injection amounts for full MS analyses, these correlations can inform pre-run peptide yield estimation to ensure the samples fall within the HeLa standard curve. Importantly, using only 10% of each sample for the initial yield estimation left sufficient material for at least one full MS analysis of all single-slide cohort samples (**Figure 1C**). The selection of injection amounts for full MS analyses balances a trade-off between analytical depth and long-term instrument performance, as well as the number of samples with sufficient material for equal injections: higher peptide loading can increase protein identifications but also necessitate more frequent maintenance. Based on these considerations, we selected a 50-ng injection for the Astral instrument. For the timsTOF analysis, we used 200 ng when sufficient material was available; otherwise, we used the remaining sample amount, but no less than 50 ng.

Across the full single-slide cohort analyses, we obtained deep proteomic coverage, identifying up to approximately 8,000 proteins (gene-centric) per sample and 10,000 proteins across the cohort (**Figure 1D–F**). The median number of proteins was 7,097 and 6,629 in the Astral and timsTOF datasets, respectively. While the analytical depth in the single-slide cohort was significantly lower than in the multi-section cohort (median 7,557 vs 7,097 and 6,980 vs 6,629 proteins in the Astral and timsTOF datasets, respectively; **Figures 1D; S1E**), higher injection amounts, as expected, increases the number of identifications (**Figure S1F,G**).

We next evaluated the biological and clinical relevance of the generated proteomic profiles. To provide biological context, we integrated publicly available protein lists of interest and clinical annotations with our MS data (**Figures 2A,E; 3D**). MS-based quantification of histological markers routinely used in pathology to distinguish adenocarcinoma (AC) from squamous cell carcinoma (SCC) enabled resolution of AC and SCC samples (**Figures 2B; S1H**). While the two histological groups were not completely separated, the strong concordance between Astral and timsTOF datasets (**Figure 2C**) suggests that the observed overlap reflects intrinsic tumor biology and marker expression heterogeneity rather than quantitative inaccuracies of the proteomics analyses.

**Figure 2.**
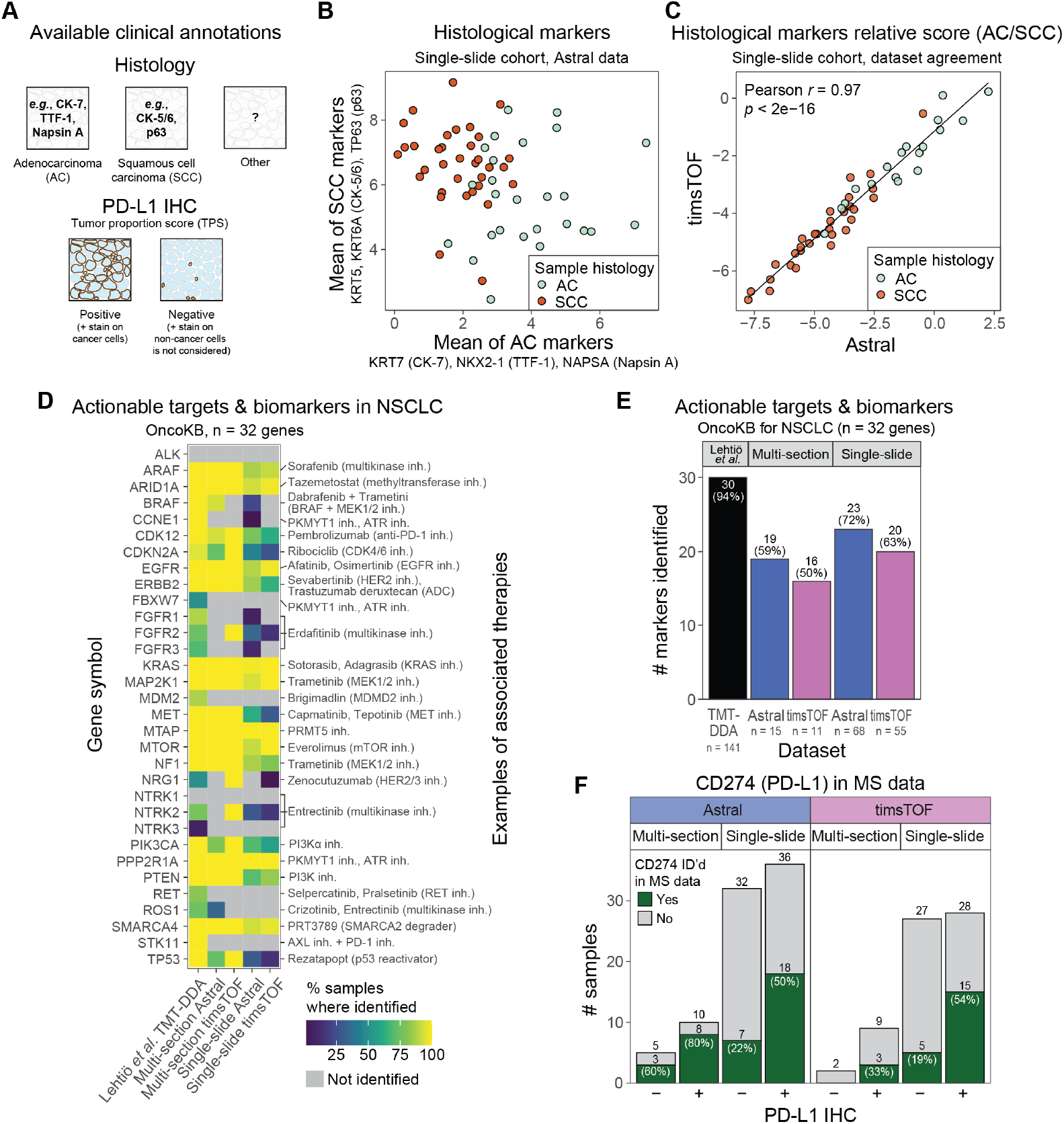
MS-based analysis of single FFPE slides provides biologically sound data and identifies clinically actionable targets. **A)** Schematic overview of the available clinical annotations used to provide context for the proteomic observations. **B)** MS-based quantitative signature of histological markers in adenocarcinoma (AC, n = 26) and squamous cell carcinoma (SCC, n = 34) samples in the single-slide cohort Astral dataset. **C)** Relative levels of histological markers (mean of adenocarcinoma markers divided by mean of squamous cell markers, as shown in panel B) in AC (n = 18) and SCC (n = 30) samples from the single-slide cohort in the Astral and timsTOF datasets. **D–E)** Identification of actionable drug targets and predictive biomarkers in non-small-cell lung cancer (NSCLC) as listed in the OncoKB database with **D)** annotation of the sample percentage where identified and **E)** summary statistics. An in-depth TMT-DDA dataset of 141 NSCLC tumors covering 13,975 proteins (Lehtiö *et al*., 2021)^5^ was used as a reference proteomics dataset. **F)** Identification of CD274 (PD-L1) in MS proteomics data stratified by PD-L1 status, as determined by tumor proportion score (TPS) assessment performed in clinical routine, where TPS > 0 is positive (+) and TPS = 0 is negative (–) PD-L1 expression.

We next assessed identification of clinically actionable drug targets and predictive biomarkers listed in the OncoKB database for lung cancer^9^. Across datasets, 63–72% of these markers were identified in the single-slide analyses, compared with 50–59% in the multi-section datasets, a difference most likely attributable to the small cohort size in the context of cancer heterogeneity (**Figure 2D,E**). We then examined MS-based identification of PD-L1 in relation to tumor PD-L1 status determined by immunohistochemistry (IHC), which is routinely used to guide immunotherapy selection in lung cancer. CD274 (PD-L1) was identified in 80% of IHC-positive cases in the multi-section Astral dataset, but in only 33–54% of positive samples in the remaining datasets (**Figure 2F**), suggesting limited sensitivity for CD274 detection in FFPE samples using the current workflows. Overall, CD274 was detected in 25–75% of samples across datasets. Notably, most identifications occurred in PD-L1 IHC-positive tumors (**Figure 2F**), whereas the remaining detections likely reflect PD-L1 expression by non-cancer cells in the tumor microenvironment (which is not captured by the IHC scoring), or tumor heterogeneity, as the PD-L1 IHC staining was performed on tissue sections distinct from those used for MS analysis.

Individual biomarkers such as PD-L1 can inform clinical decision making; however, multivariate biomarkers are increasingly recognized as required to capture the complexity of tumor biology, therapy response, and resistance mechanisms for more precise therapy selection^10^. The ability to generate data at scale for quantitative analysis of marker sets and molecular signatures is a key strength of MS-based proteomics. Here, we demonstrate signature-level analyses using the tumor inflammation signature (TIS) composed of 18 markers proposed by Ayers *et al*.^11^, which characterizes the tumor immune microenvironment (TIME), a key determinant of response to immunotherapy, (**Figure 3A,B**)^12^. Although originally defined at the mRNA-level data and now implemented in RNA-based gene expression assays; proteomics offers several advantages over transcriptomics. Proteome-level information reflects the final molecular tumor phenotype and drug target levels more accurately due to frequently low mRNA-protein correlations^5,13^. Moreover, proteins are more stable than RNA, enabling more robust profiling of archived material^14^. In the single-slide cohort, the TIS score showed strong agreement between the Astral and timsTOF datasets (**Figure 3C**). Notably, although the TIS scores were significantly higher in PD-L1-positive tumors than in PD-L1-negative, the difference was small, and the large variability in both groups highlights that TIME assessment provides information orthogonal to PD-L1 status.

**Figure 3.**
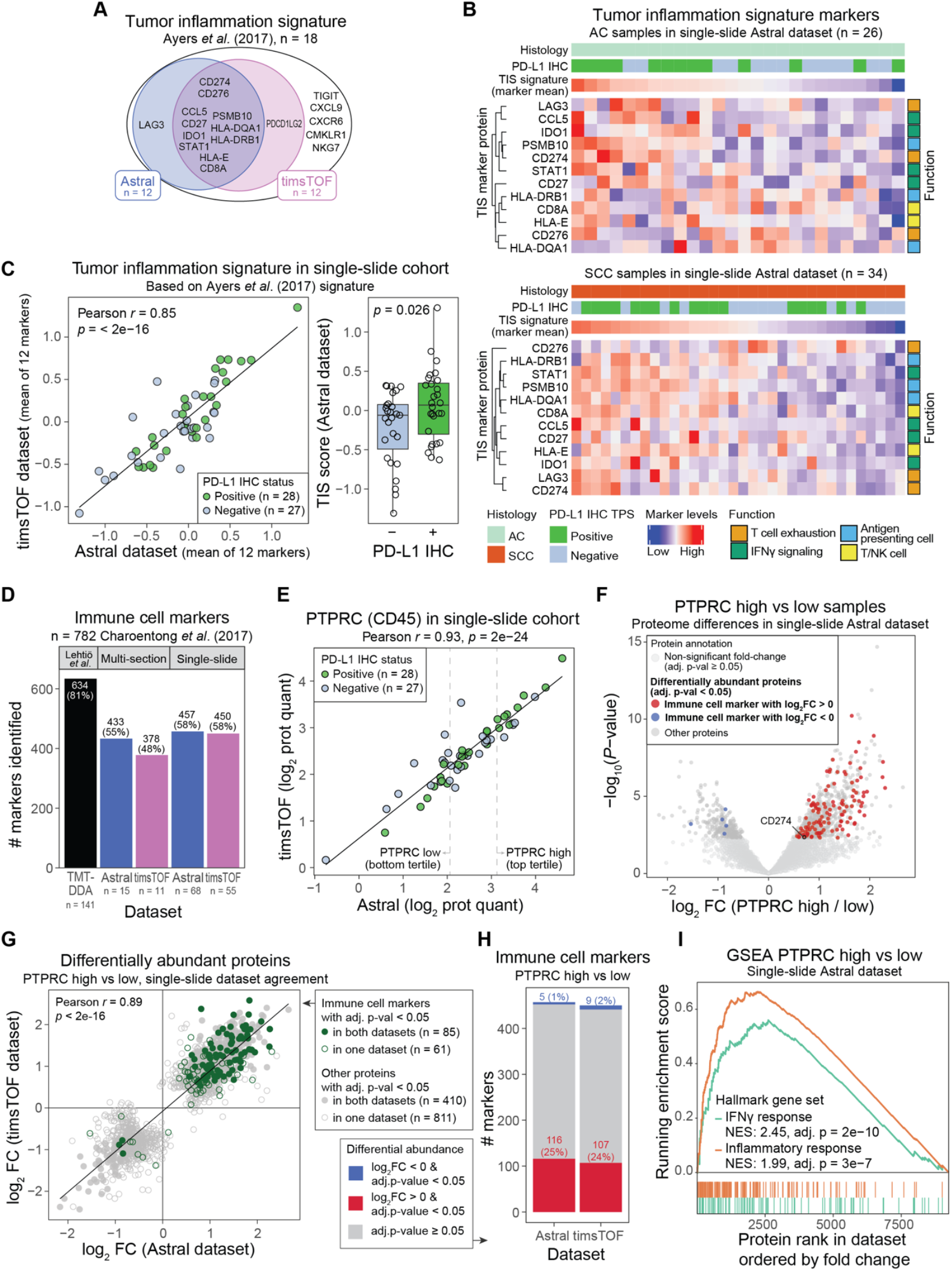
MS-based proteomics of single FFPE slides quantifies molecular signatures and provides opportunities for in-depth exploratory analyses. **A)** Markers from the tumor inflammation signature (TIS) proposed by Ayers *et al*. (2017)^11^ and identified in the single-slide cohort. **B)** Scaled levels of TIS proteins in adenocarcinoma (AC, top) and squamous cell carcinoma (SCC, bottom) samples in the single-slide Astral dataset. Columns ordered by the signature marker mean; rows clustered by hierarchical clustering using Spearman correlation distance. PD-L1 immunohistochemistry (IHC) annotation like in Figure 2. **C)** Mean of scaled TIS protein levels in single-slide Astral and timsTOF datasets (left) and stratified by PD-L1 status (right). *P* value was calculated using two-sided Welch two-sample t-test. **D)** Identification of immune cell markers published by Charoentong *et al*. (2017)^15^. The same Lehtiö *et al*. (2021)^5^ reference dataset was used as in Figure 2. **E)** Protein quantities of an immune cell marker PTPRC (CD45) in the single-slide cohort. Tertiles were defined based on the Astral dataset and included samples not analyzed in the timsTOF dataset. **F)** Differential abundance analysis using DEqMS comparing PTPRC high versus low tertiles in the single-slide cohort Astral dataset (n = 23 per group). Immune cell marker labels refer to the Charoentong *et al*. (2017) list. **G)** Comparison of differentially abundant proteins like in panel F indicated in the Astral and/or timsTOF datasets (sample groups defined using Astral data). **H)** Percentage of immune cell markers (Charoentong *et al*., 2017 list) that are differentially abundant in the PTPRC high versus low comparison in the Astral and/or timsTOF datasets. **I)** Gene-set enrichment analysis (GSEA) using ranked fold changes in PTPRC high versus low tertile comparison in the single-slide cohort Astral dataset (shown in panel F), assessing enrichment of MSigDB interferon gamma (IFNγ) and inflammatory response hallmark gene sets^16^. FC, fold change; NES, normalized enrichment score.

Beyond molecular signature quantification, in-depth proteomics enables exploratory analyses; therefore, we next examined immune-related signals at a broader scale. Using immune cell-type gene sets published by Charoentong *et al*.^15^ including 782 proteins as a reference, we detected 58% of the markers in the single-slide cohort (**Figure 3D**). We then stratified the single-slide cohort by abundance of the pan-immune cell marker PTPRC (CD45; **Figure 3E**) and compared proteomic profiles between the top and bottom tertiles (**Figure 3F–H**). In the Astral dataset, 25% of identified immune cell markers were significantly elevated in the PTPRC-high group, whereas only 1% were significantly lower. Finally, gene-set enrichment analysis (GSEA) using immune-related MSigDB hallmark gene sets^16^, as expected, revealed significant enrichment of IFNγ and inflammatory response pathways in the PTPRC-high group (**Figure 2I**).

In conclusion, our study demonstrates that state-of-the-art MS instrumentation and optimized workflows enable rapid, in-depth proteomic analysis of clinical FFPE specimens, including small biopsies and limited material on single glass slides. Although multiple sections remain advantageous for maximizing analytical depth, single-slide analyses nonetheless yield biologically and clinically informative data suitable for both targeted and exploratory applications. By supporting quantification of clinically actionable proteins, immune-related markers, and multivariate signatures, our findings highlight the value of proteomics for capturing functional tumor phenotypes beyond single-marker assessments, consistent with growing evidence that complex biomarkers are essential for precision oncology. Importantly, we show that these capabilities extend to archived FFPE biopsies derived from unresectable cases, enabling retrospective analyses of clinically annotated cohorts despite limited tissue availability. This opens opportunities for detailed proteome analysis of previously underutilized specimen collections of biopsies from late-stage patient cohorts, a patient population where new oncology drugs are typically initially tested. Our method demonstrated that clinical proteomics is a practical and powerful modality for precision medicine and cancer research.

## Methods

### Clinical sample collection

The multi-section FFPE cohort comprised of 15 samples collected from lung cancer patients at the Karolinska University Hospital in Solna, Sweden. The surgical samples originated from resected lung tumors during routine treatment of patients. The biopsy samples (core needle or forceps) were collected during diagnostic procedures for suspected lung cancer patients. The study was approved by the Regional Ethical Review Board in Uppsala, Sweden (registration no. 2021-01931 and 2024-04958-02). Informed consent was obtained from all the patients. The FFPE sample preservation was performed as part of the clinical routine. The samples were taken for research only after the routine diagnostics was performed and there was sufficient material left. Macrodissection was performed to minimize the amount of non-tumor material. Five consecutive cuts of 5 μm each were then taken for and shortly before proteomic analysis.

The single-slide FFPE cohort comprised of 68 samples collected from lung cancer patients at the Oslo University Hospital in Oslo, Norway. The biopsy samples (core needle, forceps or cryobiopsy) were collected as part of the DART trial approved by the Regional Ethical Committee for Medical Health Research Ethics, REK South-East in Oslo, Norway (reference no. 48655). Informed consent was obtained from all the patients. The FFPE sample preservation was performed as part of the clinical routine with 4-μm tissue sections mounted on glass slides. The glass slides were initially prepared for other analyses, leftover slides were stored for several years, after which they were used for proteomic analyses.

The meta data for the samples, including histological and PD-L1 status annotations, peptide yields, generated datasets and the number of identified proteins is provided in **Supplementary Data**.

### Sample preparation

For the single-slide cohort, the slides were pre-washed with 0.01% n-Dodecyl-B-D-Maltoside (DDM) by dipping and shaking the slide in the solution for approximately 10 s, rinsing in water twice and drying. The FFPE material was then scraped off using a sterile scalpel into Eppendorf tubes.

All samples were prepared for proteomics analysis using the FenoPrep™ kit (Cat# 532001, FenoMark Diagnostics AB, Sweden) per the manufacturer’s instructions. Briefly, the lysis buffer was added to the samples, heated under shaking (95 °C, 400 rpm, 30 min), after which the sample tubes were placed in the VialTweeter Ultrasonic processor (Hielscher Ultrasonics GmbH, Germany; settings: amplitude 100%, pulsation mode 100%, 10 cycles of 60 s on/30 s off). The samples were then heated under shaking (95 °C, 400 rpm, 30 min). The samples were then placed in the ultrasonic processor for 5 more cycles. The samples were centrifuged (13,000 × g, 10 min) and the supernatant was transferred for further processing. The proteins were alkylated. Thereafter, on-bead digestion using Lys-C and trypsin and clean-up using magnetic beads were performed. For the multi-section cohort, the peptide concentration was measured using a microBCA assay with HeLa digest as the standard. For the single-slide cohort, MS-based peptide yield estimation was performed (see next section).

### MS-based peptide yield estimation using timsTOF HT

A timsTOF HT mass spectrometer (Bruker) connected to an EVOSEP LC system via the CaptiveSpray 2 source was used for DIA-PASEF LC-MS/MS analysis. Each sample was analyzed after transfer of 10% peptide solution volume on Evotips following the manufacturer’s instructions (except for one sample where 1% of the volume was used, loading amounts are provided in the supplementary materials). The peptides were separated on a PepSep C18 column (Bruker, 8 cm x 150 µm, 1.5 µm), using the 60SPD fixed method. For the dia-PASEF analysis the window scheme was calculated using the py_diAID tool (https://github.com/MannLabs/pydiaid). The capillary voltage was set at 1500 V, stepping collision energy at 32, 40, 50 eV, dia-PASEF scan range 100–1700 m/z in positive mode, and IMS service ramp time of 100 ms.

### DIA analysis using Orbitrap Astral

For the multi-section cohort, an Orbitrap Astral coupled to a Vanquish NEO System (Thermo Fisher Scientific) was used. The injection volume was 2 μl for each sample containing 500 ng of peptides. The samples were trapped on a C18 guard-desalting column (Acclaim PepMap NEO, 300μm x 5 mm, nanoViper, C18, 5 µm, 100 Å) and separated on a 25 cm long C18 column (25 cm Aurora Ultimate XT column, C18, 1.7 μm, 75 μm x 25 cm). The nanocapillary solvent A was 0.1% formic acid in water; and solvent B was 0.1% formic acid in acetonitrile. At a constant flow of 0.25 μl/min, the curved gradient went from 6% B up to 40% B in 32 min, followed by a steep increase to 100% B in 5 min.

For the single-slide cohort, an Orbitrap Astral System (Thermo Fisher Scientific) coupled to an EVOSEP LC system was used. Each sample was analyzed after the transfer of 50 ng of peptides on Evotips following the manufacturer’s instructions. The peptides were separated on a PepSep C18 column (Bruker, 15 cm x 75 µm, 1.5 µm), using the 30SPD fixed method with a total method duration of 44 min.

The standard MS parameters for DIA acquisition were set as follows: a spray voltage of 2000 V, no sheath or auxiliary gas flow, and a heated capillary maintained at 280 °C. Full-scan mass spectra were acquired with a scan range of 380–980 m/z, an AGC target value of 300%, maximum injection time (IT) of 10 ms, and a resolution of 240,000. The DIA windows were set to 2 m/z in a precursor mass range of 380–980 m/z and subjected to fragmentation (nCE: 25%) with a scan range of 145–1450 m/z, maximum IT of 3 ms, AGC target value of 500%.

### DIA analysis using timsTOF HT

A timsTOF HT mass spectrometer (Bruker) connected to an EVOSEP LC system via the CaptiveSpray 2 source was used for DIA-PASEF LC-MS/MS analysis.

For the multi-section cohort, 11 samples were analyzed after transfer of 500 ng of peptides on Evotips following the manufacturer’s instructions. The peptides were separated on a C18 IonOpticks column (particle size 1.6 µm, 75 uM µm ID, 15 cm length) at a flow rate of 0.25 µl/min; solvent A (0.1% formic acid in water), solvent B (0.1% formic acid in acetonitrile), using the 30SPD fixed method with a total method duration of 44 min.

For the single-slide cohort, 56 samples sample was analyzed after transfer of 50–200 ng of peptides on Evotips following the manufacturer’s instructions (loading amounts are provided in the **Supplementary Data**). The peptides were separated on a PepSep C18 column (Bruker, 15 cm x 75 µm, 1.5 µm), using the 30SPD fixed method with a total method duration of 44 min.

For the dia-PASEF analysis the window scheme was calculated using the py_diAID tool (https://github.com/MannLabs/pydiaid). The capillary voltage was set at 1500 V, stepping collision energy at 32, 40, 50 eV, dia-PASEF scan range 100–1700 m/z in positive mode, and IMS service ramp time of 100 ms.

One injection in the single-slide cohort failed, resulting in the final dataset of 55 samples.

### DIA analysis using Orbitrap Exploris 480

An Orbitrap Exploris 480 coupled to a Dionex UltiMate™ 3000 RSLCnano System was used System (Thermo Fisher Scientific). The injection volume was 2 µl for each sample containing 2 µl. Samples were trapped on a C18 guard-desalting column (Acclaim PepMap 100, 75 μm x 2 cm, nanoViper, C18, 5 µm, 100 Å) and separated on a 25-cm long C18 column (25 cm Aurora Ultimate XT column, C18, 1.7 μm, 75 μm x 25 cm). The nanocapillary solvent A was 0.1% formic acid in water; and solvent B was 0.1% formic acid in acetonitrile. At a constant flow of 0.25 μl/min, the curved gradient went from 6% B up to 40% B in 120 min followed by a steep increase to 100% B in 5 min. The standard MS parameters for DIA acquisition were set as follows: a spray voltage of 1900 V, no sheath or auxiliary gas flow, and a heated capillary maintained at 275 °C. Full-scan mass spectra were acquired with a scan range of 375–1175 m/z, an AGC target value of 100%, maximum injection time (IT) of 100 ms, and a resolution of 120,000. The DIA windows were set to 25 m/z in a precursor mass range of 375–1175 m/z and subjected to fragmentation (nCE: 28%) by HCD with a scan range of 145–1450 m/z, maximum IT of 100 ms, AGC target value of 100%, and a resolution of 15000.

### DIA data search and preprocessing

DIA data was analyzed in Spectronaut (Biognosys, v.20) using directDIA mode and searched against ENSEMBL protein database (GRCh38.pep.all). The peptide quantities were summarized into proteins (gene-centric) using the MaxLFQ algorithm and only gene-specific peptides. The data was normalized using local normalization. Both imputed (global imputing) and non-imputed data were exported. Additionally, precursor-level non-imputed and non-normalized data, as well as the total ion chromatograms were exported. Non-imputed data was used for identification analyses, while imputed exports were used as quantitative data. Total ion chromatograms and precursor-level exports were used for MS signal calculations; additionally, precursor-level data was used to assess the contribution of high-abundance proteins.

### Downstream bioinformatics analysis

All downstream bioinformatics analyses were performed in R (v. 4.5.2). To calculate the sample-wise normalized protein quantities, the MS2 area values exported from Spectronaut were divided by the median of all quantification in the respective sample; thereafter the quantities were log_2_ transformed. For high abundance protein analysis, precursor-level MS1 data was exported. For the total ion chromatogram area under the curve (TIC AUC) analysis, the TIC chart data was exported from Spectronaut. Thereafter, the AUC was calculated using the AUC() function (trapezoid method) from the DescTools R package (v. 0.99.60). The differential abundance analysis was performed using the DEqMS package^17^ (v. 1.26.0) using imputed data and median precursor counts per protein. *P* values were adjusted using the BH method. The gene set enrichment analysis (GSEA) was performed using the GSEA() function from the clusterProfiler R package^18^ (v. 4.16.0) based on the ranked fold-change values derived from the DEqMS analysis. The hallmark gene sets originated from the MSigDB database^16^ (msigdbr R package v. 25.1.1). All the used statistical tests and the corresponding sample sizes are specified in the figures and figure legends.

### Public domain datasets

The Charoentong *et al*. immune cell markers were taken from the supplementary tables of the original publication^15^. The IFN gamma response and inflammatory response hallmark gene sets originated from the MSigDB^16^.

## Data availability

MS proteomics data have been deposited on the ProteomeXchange Consortium via the PRIDE partner repository. The datasets will be made publicly available upon publication of the peer-reviewed manuscript.

## Acknowledgements

We thank FenoMark Diagnostics for providing the FenoPrep™ kit for sample preparation. This study was funded by Cancerfonden (23 2819 Pj, 25 4349 Pj), Vetenskapsrådet (ERA PerMed 2021-00754, 2024-03294), Radiumhemmets Forskningsfonder (244203), Karolinska Institutet (2023-01372), Stockholm County council (ALF FoUI-1000396), Vinnova (ProMS 2024-01137), EU DART (965397), EU Rescuer (847912). The single-slide cohort originated from the DART trial, which was a researcher-initiated study and funded by grants from the South-Eastern Norway Regional Health Authority (grant no: 2019119 and 26011) and AstraZeneca. The funder was not involved in the study design, collection, analysis, interpretation of data, the writing of this article or the decision to submit it for publication.

## Author contributions

The study was conceptualized by OB, LMO, and JL. The study was supervised by LMO. Clinical samples were collected and annotated by IS, VDH, and ÅH. Sample preparation was performed by NEÖ, GM, and MN. Mass spectrometry data generation was performed by GM and NEÖ. Data analysis was performed by OB. The manuscript was written by OB. All authors approved the final version of the manuscript.

## Ethics declarations

LMO, JL, and GM are shareholders of FenoMark Diagnostics that provides proteomics solutions to medical professionals, including a kit that was used in this study. ÅH has participated in meetings (talks/advise/discussions) arranged by Abbvie, Takeda, AstraZeneca, Roche, Pfizer, Janssen, Eli Lilly, BMS, PierreFabre, Bayer, MSD, Novartis, Merck, Sanofi, Medicover, BeOne. All fees were paid to OUS. ÅH has received support for research projects and studies from Roche, AstraZeneca, Novartis, Incyte, EliLilly, Bristol-Myers Squibb, Ultimovacs, Merck, GSK, Medicover, BeOne, Johnson & Johnson. VDH has participated in lectures organized by Bristol-Myers Squibb, MSD, AstraZeneca, Johnson&Johnson, BeOne, and Regeneron; and has participated in advisory boards of MSD, AstraZeneca, Bristol-Myers Squibb, BeOne, GSK, Immedica, Merck, and Novartis.

## Supplementary materials

**Supplementary Table 1.** Cohort characteristics.

|  | **Multi-section cohort**<br>**(N = 15)** | **Single-slide cohort**<br>**(N = 68)** |
| --- | --- | --- |
| Median age, years (range) | 75 (48–85) | 68 (36–85) |
| Sex |  |  |
| Male | 7 (47%) | 39 (57%) |
| Female | 8 (53%) | 29 (43%) |
| Histology |  |  |
| Adenocarcinoma | 15 (100%) | 26 (38%) |
| Squamous cell carcinoma | 0 | 34 (50%) |
| Other or NA | 0 | 8 (12%) |
| Stage |  |  |
| I | 5 (33%) | 0 |
| II | 3 (20%) | 1 (1%) |
| III | 3 (20%) | 65 (96%) |
| IV | 4 (27%) | 0 |
| Unknown | 0 | 2 (3%) |
| PD-L1 status |  |  |
| Negative | 5 (33%) | 32 (47%) |
| Positive | 10 (67%) | 36 (53%) |
| TPS 1–49 | 5 (33%) | NA |
| TPS ≥50 | 5 (33%) | NA |
| Sample type |  |  |
| Surgical | 7 (47%) | 0 |
| Biopsy | 8 (53%) | 68 (100%) |
| Core biopsy | 5 (33%) | 32 (47%) |
| Forceps biopsy | 3 (20%) | 32 (47%) |
| Cryobiopsy | 0 | 4 (6%) |
| MS data |  |  |
| Astral | 15 (100%) | 68 (100%) |
| timsTOF | 11 (73%) | 55 (81%) |
| Exploris | 11 (73%) | NA |

**Supplementary Figure S1.**
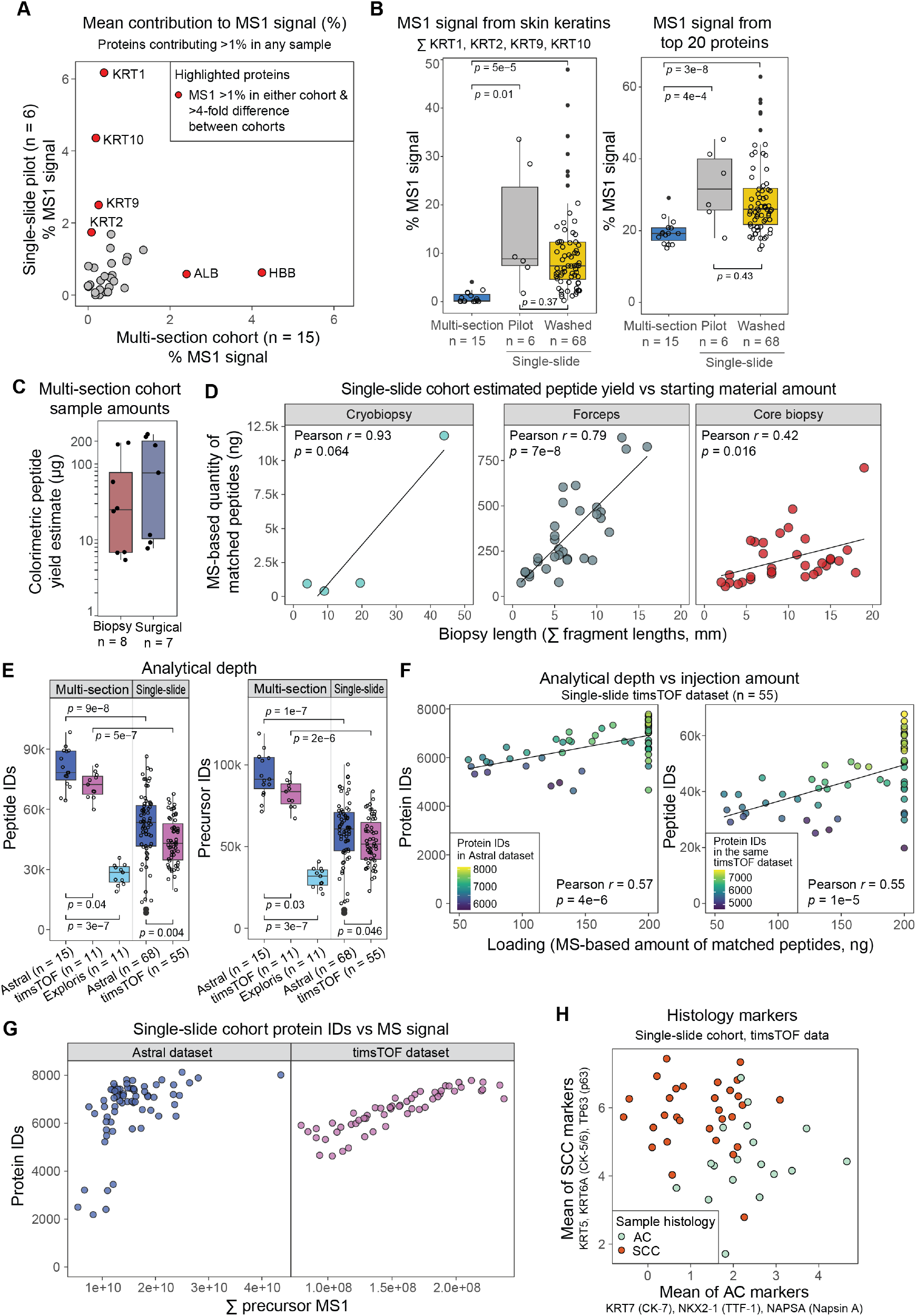
Analytical characteristics of the MS-based analysis of formalin-fixed, paraffin embedded (FFPE) clinical tumor samples (related to Figures 1 and 2). **A)** High abundance proteins (defined as contributing more than 1% of total MS1 signal in any sample, n = 31) and their mean contribution to the total MS1 signal across the multi-section and single-slide cohorts. **B)** Percentage of MS1 signal attributed to high-abundance proteins: four skin keratins (left) and top 20 proteins by signal (right) in the multi-section cohort, pilot analysis of the single-slide cohort, and the full analysis of the single-slide cohort, including a slide washing step. *P* values were calculated using Wilcoxon rank-sum test. The number of samples in each dataset is indicated in the figure. **C)** Peptide yield per sample in the multi-section cohort (n = 15) as determined using a colorimetric assay, grouped by sample type: biopsy (core or forceps, n = 8) and surgical material (n = 7). **D)** Peptide yields (as determined using the MS-based approach using HeLa standards) in the single-slide cohort in relation to the size of the biopsy on the slide, grouped by biopsy type: cryobiopsy (n = 4), forceps biopsy (n = 32) and core biopsy (n = 32). **E)** Number of peptides (left) and precursors (right) identified per sample in MS-based analyses of the multi-section and single-slide cohorts. *P* values were calculated using Wilcoxon rank-sum test. The number of samples in each dataset is indicated in the figure. **F)** Number of proteins (gene-centric, left) and peptides (right) identified per sample in the single-slide cohort timsTOF dataset, in relation to the injected sample amount. **G)** Number of proteins (gene-centric) identified per sample in the single-slide cohort in relation to the total MS signal expressed as the sum of precursor MS1 signal, shown for the the Astral (left) and timsTOF (right) datasets. **H)** Histology markers in adenocarcinoma (AC) and squamous cell carcinoma (SCC) samples (n = 48) in the single-slide cohort timsTOF dataset.

